# Exogenous Recombinant N-Acetylgalactosamine-4-Sulfatase (Arylsulfatase B; ARSB) Inhibits Progression of B16F10 Cutaneous Melanomas and Modulates Cell Signaling

**DOI:** 10.1101/2023.05.30.542851

**Authors:** Sumit Bhattacharyya, Insug O-Sullivan, Jieqi Tu, Zhengjia Chen, Joanne K. Tobacman

## Abstract

In the syngeneic, subcutaneous B16F10 mouse model of malignant melanoma, treatment with exogenous ARSB markedly reduced tumor size and extended survival. *In vivo* experiments showed that local treatment with exogenous N-acetylgalactosamine-4-sulfatase (Arylsulfatase B; ARSB) led to reduced tumor growth over time (p<0.0001) and improved the probability of survival up to 21 days (p=0.0391). Tumor tissue from the treated mice had lower chondroitin 4-sulfate (C4S) content and lower sulfotransferase activity. The free galectin-3 declined, and the SHP2 activity increased, due to altered binding with chondroitin 4-sulfate. These changes induced effects on transcription, which were mediated by Sp1, phospho-ERK1/2, and phospho-p38 MAPK. Reduced mRNA expression of chondroitin sulfate proteoglycan 4 (CSPG4), chondroitin sulfotransferase 15 (N-acetylgalactosamine 4-sulfate 6-O-sulfotransferase), and matrix metalloproteinases 2 and 9 resulted. Experiments in the human melanoma cell line A375 demonstrated similar responses to exogenous ARSB as in the tumors, and inverse effects followed RNA silencing. ARSB, which removes the 4-sulfate group at the non-reducing end of C4S, acts as a tumor suppressor, and treatment with exogenous ARSB impacts on vital cell signaling and reduces the expression of critical genes associated with melanoma progression.

**Highlights:**

Exogenous ARSB reduced tumor size and increased survival
Chondroitin 4-sulfate increased, leading to increased free galectin-3
mRNA expression of CSPG4 and CHST15 declined following ARSB treatment
mRNA expression of MMP9 and pro-MMP2 declined following ARSB treatment
Active SHP2 increased, leading to declines in phospho-ERK1/2 and phospho-p38 MAPK

## 1.0 Introduction

**I**n cultured, human malignant melanoma cells, the expression of the enzyme N-acetylgalactosamine-4-sulfatase (Arylsulfatase B; ARSB) declined with increasing invasiveness of a series of seven human melanoma cell lines [1]. As ARSB declined, the chondroitin 4-sulfate (C4S) content increased, since ARSB removes the 4-sulfate group at the non-reducing end of the C4S or dermatan sulfate chain and is required for the degradation of these sulfated glycosaminoglycans [2–6]. Defective ARSB in the congenital disorder Mucopolysaccharidosis (MPS) VI leads to accumulation of C4S and dermatan sulfate and systemic manifestations, characterized by skeletal abnormalities, short stature, hepatomegaly, and reduced life expectancy [7]. Early treatment with recombinant ARSB can ameliorate some of the disease pathobiology [7].

In cell-based studies, effects of decline in ARSB on cellular mediators, which either bind directly to C4S or are regulated by C4S-binding partners, have been identified [8]. Proteins which bind differentially to C4S, due to effects of ARSB on removal of the sulfate group at the non-reducing end, include galectin-3, non-receptor tyrosine phosphatase SHP2 (PTPN11), kininogen, IL-8, and BMP4 [8]. Free galectin-3 increases when ARSB is inhibited and C4S is more sulfated [1,9–7,11]. Unbound to C4S, galectin-3 interacts with transcription factors, including Sp1 and AP-1, to enhance expression of proteins engaged in vital cell functions, such as Wnt9A and HIF-1α, and proteins which form chondroitin sulfate proteoglycans, such as versican and chondroitin sulfate proteoglycan 4 (CSPG4) [1,9–11].

Prior work detected increased expression of CSPG4, also known as melanoma-associated chondroitin sulfate proteoglycan (MCSP) or neuron-glial antigen 2 (NG2), when ARSB activity was reduced. When C4S increased, bound galectin-3 declined, free galectin-3 increased, and galectin-3 with Sp1 enhanced CSPG4 promoter activation [1]. CSPG4 has been recognized for decades as a biomarker of melanoma progression [12,13], and novel treatments directed at inhibition of CSPG4 are under development [14–17]. In this project, following treatment with recombinant human ARSB in the syngeneic B16F10 mouse model of subcutaneous melanoma and in the human A375 melanoma cell line declined significantly, expression of CSPG4 declined, suggesting another approach to inhibit CSPG4.

In addition to impact on CSPG4 expression, treatment of the mouse melanomas with exogenous ARSB increased the activity of SHP2, attributable to less binding of SHP2 with C4S following ARSB, as previously [1,18–21]. Increased activity of SHP2 led to decline in phosphorylation and activity of p38-MAPK and ERK1/2, and reduced the expression of the metalloproteinases MMP-2 and MMP-9. MMP-2 and MMP-9 are Type IV collagenases involved in the remodeling of the extracellular matrix. They are well-recognized as facilitators of tumor invasiveness and metastatic proclivity, and anti-cancer therapy directed at reducing their expression by use of tissue inhibitors of metalloproteinases (TIMPs) has been undertaken previously [22–31]. The impact of exogenous ARSB on reducing their expression suggests another approach to inhibit the impact of the MMPs on melanoma progression.

Malignant melanoma continues to increase in prevalence with associated high morbidity and mortality. Although responses to immunotherapy by checkpoint inhibitors are highly effective for some individuals, other melanoma patients are unresponsive or become refractory. With this background and the ongoing need for better therapies, the impact of rhARSB on the progression of cutaneous melanomas in the syngeneic mouse melanoma B16F10 model was investigated. Exogenous ARSB has been used safely and effectively for almost 20 years in MPS VI [7], with replacement dose of 1.0 mg/kg IV weekly. Bioactive ARSB at a dose of 0.1-0.4 mg/kg subcutaneously was tested at intervals of 2-7 days in the studies in this report.

## 2.0 Materials and Methods

### 2.1 B16F10 syngeneic mouse model

The B16F10 melanoma cell line was purchased from ATCC (Manassas, VA, USA). Cells were cultured in Dulbecco’s modified Eagle medium supplemented with 10% fetal bovine serum (FBS), 1% penicillin-streptomycin. Cells were screened for pathogens by IDEXX BioAnalytics (Columbia, MO). The cells were maintained at 37°C in a humidified, 5% CO_2_ environment with media exchange every 2 days. Confluent cells in T-25 flasks were harvested by EDTA-trypsin, and sub-cultured. Eight-week-old female C57BL/6J mice (n = 40) were purchased (Jackson Laboratories, Bar Harbor, Maine, USA) and housed in the Veterinary Medicine Unit at the Jesse Brown VA Medical Center (Chicago, IL, USA). Principles of laboratory animal care were followed, and all procedures were approved by the AALAC accredited Animal Care Committee. Mice were fed a standard diet and maintained in groups of three in a cage with routine light–dark cycles. The animals were divided into five groups, control tumor group (n=12), four recombinant ARSB-treated groups (n=28 total) and ARSB injection control group with no tumor (n=3). Mice were inoculated subcutaneously with 2.5 x 10^5^ B16F10 cells in 100 μl of normal saline. Body weight and tumor volume were measured using calipers, and the volume was expressed in cm^3^ [0.5 × L × W^2^] (L=long diameter; W=short diameter of the tumor) [32].

Recombinant ARSB treatment was started 48 hours post tumor inoculation. Recombinant bioactive ARSB (R&D Systems, Minneapolis, MN, USA) in normal saline (n=12) was injected subcutaneously around the tumor with 25-gauge needle. In three of the treatment groups, 0.1 mg/kg BW (n=5), 0.2 mg/kg BW (n=11), or 0.4mg/kg BW (n=6) of rhARSB was injected on days 2, 7, and 14. A fourth treated group (n=6) received 0.2 mg/kg BW of rhARSB more frequently, on days 2, 7, 11, 15, and 19.

A375 human melanoma cells (CRL-1619, ATCC) were grown in DMEM with 10% FBS to ∼70% confluence, then silenced using ARSB siRNA and galectin-3 siRNA, in some experiments, as previously [1]. Media exchange occurred after 24h and cell treatments were introduced 24h after silencing began. Treatments were for 24h, unless indicated otherwise, and included: PHPS1 (phenylhydrazonopyrazolone sulfonate (30 μM; Sigma-Aldrich, St. Louis, MO, USA), a chemical inhibitor of SHP2; mithramycin (250 nM; Sigma), an inhibitor of Sp1 binding to DNA; SB203580 SB203580, a highly specific p38 MAPK inhibitor (10 µM; EMD Millipore, Billerica, MA); and ERK activation inhibitor peptide 1, cell-permeable (10 μM; Sigma) [1,9,12,13]. Some cell preparations were treated with rhARSB (1 ng/ml x 24h) or silenced by ARSB siRNA, as previously [1]. Cells were harvested and frozen at -80°C for subsequent analysis.

### 2.2 mRNA expression of CHST15, CSPG4, pro-MMP2, and MMP-9

Total RNA was prepared from treated and control tissues using RNeasy Mini Kit (Qiagen, Germantown, MD). Equal amounts of purified RNAs from the control and treated tissues were reverse-transcribed and amplified using Brilliant SYBR Green QRT-PCR Master Mix (Bio-Rad, Hercules, CA). Human β-actin was used as an internal control. QRT-PCR was performed using the following specific primers:

Mouse CHST15 (NM_029935.6) left: 5’-TATGCCCAGCGTAGAGAAGG-3’ and right: 5’- ACCAGCCAGAACCAAAAACA-3’;

Mouse CSPG4 (NM_139001.2) left: 5’-TACCTGAGCACTGACCCACA-3’ and right: 5’- TTCCCTCTTCCTCCTCTTCC-3’;

Mouse MMP2 (NM_008610) left: 5’-CAGTGATGGCTTCCTCTGGT- 3’ and right: 5’– GGTDATAGTCCTCGGTGGTG-3’;

Mouse pro-MMP9 (NM_013599.5) left: 5’-GACTACGATAAGGACGGCAAA-3’ and right: 5’- AGGGCAGAAGCCATACAGTTT-3’.

Human CHST15 (NM_015892) forward: 5′-ACTGAAGGGAACGAAAACTGG-3′ and reverse: 5′- CCGTAATGGAAAGGTGATGAG-3′;

Human CSPG4 (NM_001897) left: 5’̒-CTTCAACTACAGGGCACAAGG-3’ and right: 5’- AGGACATTGGTGAGGACAGG-3’;

Human pro-MMP2 (NM_004530) left: 5’-AGTGGATGATGCCTTTGCTC-3’ and right: 5’- GAGTCCGTCCTTACCGTCAA-3’;

Human MMP9 (NM_004994.2) left: 5’-GTCTTCCCCTTCACTTTCCTG-3’ and right: 5’- TCAGTGAAGCG GTACATAGGG-3’.

Cycle threshold (Ct) was determined during the exponential phase of amplification, as previously [9]. Fold changes in expression were determined from the differences between the Ct values using the standard procedure [9], and the expression relative to the untreated control was determined

### 2.3 Measurement of total sulfated glycosaminoglycans, total chondroitin sulfate, and chondroitin 4-sulfate

Total sulfated glycosaminoglycans (sGAG) were measured in the cell extracts by sulfated GAG assay (Blyscan^™^, Biocolor Ltd, Newtownabbey, Northern Ireland) per manufacturer’s instructions [33]. Chondroitin sulfate (CS) and chondroitin 4-sulfate in the samples were determined following immunoprecipitation with CS antibody (CS-56, Abcam, Waltham, MA, USA), or C4S antibody LY111 (TCI Chemicals, Portland, OR, USA). Dynabeads (Life Technologies, Carlsbad, CA, USA) were coated with the specific CS or C4S antibody, and beads were mixed with samples, incubated, and immunoprecipitated. Immunoprecipitated CS or C4S molecules were eluted and subjected to the Blyscan sulfated GAG assay, as above.

### 2.4 Total sulfotransferase activity

Total sulfotransferase activity was determined using the Universal Sulfotransferase Activity kit (R&D) as per manufacturer’s recommendations. Activities were normalized with total cellular protein and expressed as percentage of control.

### 2.5 Silencing of ARSB and galectin-3 by siRNA

Small interfering RNA was obtained to knockdown the expression of ARSB and galectin-3 (Qiagen, Germantown, MD). Effects on mRNA were determined by QRT-PCR. The siRNA sequences for ARSB (NM_000046) silencing were: sense 5′- GGGUAUGGUCUCUAGGCA - 3′ and antisense: 5′- UUGCCUAGAGACCAUACCC - 3′. The sequence of the DNA template for human galectin-3 silencing (Hs_ LGALS3_9) was: 5΄ - ATGATGTTGCCTTCCACTTTA - 3′. Cells were grown to ∼60% confluence, then silenced by adding 0.6 μl of 20 μM siRNA (150 ng), mixed with 100 μl of serum-free medium and 12 μl of HiPerfect Transfection Reagent (Qiagen, Germantown, MD, USA). Effectiveness of ARSB silencing was confirmed by activity assay, with decline of ∼90%. Galectin-3 silencing was confirmed by galectin-3 ELISA, with galectin-3 protein declining ∼90%.

### 2.6 Phospho-ERK1/2, phospho-p38 MAPK, and galectin-3 measurements

Phospho-ERK1(T202/Y204)/ERK2(T185/Y187), phospho-p38, SHP2 activity, and galectin-3 in the tumor extracts were measured by specific DuoSet sandwich ELISA kit or ELISA assay (R&D). Cell lysates were prepared from control and treated cells in cell lysis buffer (Cell Signaling, Danvers, MA, USA) with protease and phosphatase inhibitors (Halt Protease and Phosphatase Inhibitor Cocktail, Thermo Scientific, Pittsburgh, PA, USA). Briefly, samples and standards were added to the wells of the microtiter plate precoated with a capture antibody. Phospho-ERK, phospho-SHP2, or phospho-p38, or galectin-3 in the lysates was captured by the coated antibody on the plate and detected with a specific, biotinylated second antibody. Streptavidin-HRP and hydrogen peroxide/tetramethylbenzidine substrate were used to develop color proportional to the bound HRP activity. The reaction was stopped, and the optical density of the color was read at 450 nm in a plate reader (FLUOstar, BMG, Cary, NC, USA).

### 2.7 Oligonucleotide-based ELISA for nuclear Sp1

Oligonucleotide-binding assay (TransAM kit, Active Motif, Carlsbad, CA, USA) was used to detect nuclear Sp1 in the tumor tissue. Nuclear extracts were prepared using a nuclear extract preparation kit (Active Motif) and were added to the wells of a 96-well microtiter plate, precoated with the Sp1 consensus oligonucleotide sequence 5′-GGGGCGGGG-3′. Competition for Sp1 nuclear protein binding to the consensus oligonucleotide was performed using a Sp1-mutated oligonucleotide and an Sp1 WT consensus sequence oligonucleotide, prior to detection of bound Sp1 by Sp1 antibody. Sample values were normalized by total protein and expressed as percent of untreated control.

### 2.8 Statistical analysis

A total of 40 mice were included in the study. The tumor volumes were measured at several time points (14, 15, 16, 17, 18, 19, 20, 21, 24, 27, 29, and 31 days) and treated as continuous outcome. The mean longitudinal tumor volumes of 5 different treatment groups (group 1 = Vehicle, group 2 = 0.1 mg dose, group 3 = 0.2 mg dose, group 4 = 0.4 mg dose, and group 5 = 0.2 mg dose with high frequency) at different measurement time points were first graphically presented (**Fig. 1C).** The p-values for the pairwise comparisons of tumor volume over the followup time between each pair of groups were presented (**Fig. 1D**). A mixed effect model was then used to test: 1) whether there were any significant differences in the longitudinal tumor volume across these 5 different treatment groups; 2) whether there was any significant growth trend of tumor volume over time, and 3) pairwise group comparisons in the longitudinal tumor volumes. Survival analyses were performed using the Kaplan-Meier method, Log-rank test, and Cox proportional hazard model to investigate: 1) the Product-Limit (PL) estimates of survival probability at each time point with 95% confidence interval; 2) whether there was significant difference in the survival functions across 5 groups; 3) pairwise group comparisons with group 1 in the survival functions. The significance levels were set at 0.05 for all tests. The SAS statistical package version 9.4 (SAS Institute, Inc., Cary, North Carolina) and R version 4.2.3 were used for data managements and analyses.

**FIG. 1.**
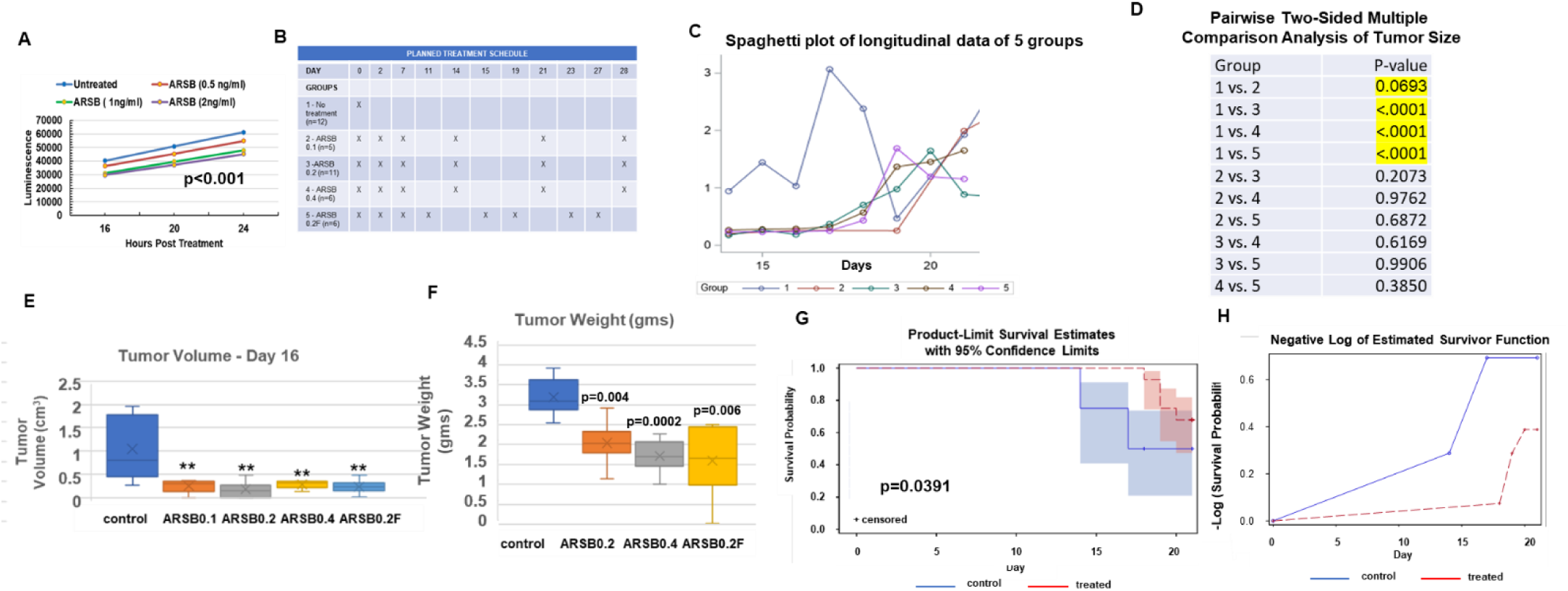
Exogenous ARSB reduces tumor size and increases survival in B16F10 subcutaneous tumor model. **A.** BrdU incorporation was less in the human melanoma cell line A375 following treatment with exogenous ARSB at concentrations of 0.5 ng/ml, 1 ng/ml and 2 ng/ml (p<0.001 for each at 24 h, two-sample t-test, two-tailed, unequal variance). **B.** The schedule of treatment and the five groups, including control (Group 1), 0.1 ng/ml (Group 2), 0.2 ng/ml (Group 3), 0.4 ng/ml (Group 4), and 0.2 ng/ml more frequent administration of ARSB (Group 5, labeled ARSB 0.2F) are indicated. **C.** The mean longitudinal tumor volumes of the 5 different groups is presented in a spaghetti plot at different measurement time points to day 21. Tumor volume was significantly less in the ARSB-treated mice than the untreated control (p<0.0001, mixed effect model). **D.** Tumor volume was smaller following treatment with rhARSB in the mice treated with exogenous ARSB (Groups 3, 4, and 5; p<0.0001, mixed effect model). Tumor volume in mice treated with 0.1 mg/kg exogenous ARSB was not significantly smaller than in controls (p=0.07). **E.** Representative data from day 16 show that average tumor volume in the untreated mice was significantly greater than in the treated mice. **F.** Tumor weight at the time of death, either spontaneous death or euthanasia, was significantly greater in the untreated control than in the groups treated with at least 0.2 mg/kg of exogenous ARSB. **G.** Survival analysis at 21 days indicated that the treated mice (including those treated with 0.1 mg/kg) had longer survival than controls (p=0.0391, log rank test), as shown in the Product-Limit Survival Estimates with 95% confidence limits. **H.** The Negative-Log of Estimated Survivor Function (F) shows improved survival in the treated mice, compared to control. [ARSB=arylsulfatase B; group 1=untreated control; group 2= ARSB 0.1 mg/kg; group 3=ARSB 0.2=0.2 mg/kg; group 4=ARSB 0.4=0.4 mg/kg; group 5=ARSB 0.2F=02 mg/kg more frequently]

Measurements from tumor samples are the mean ± SD of at least six independent experiments. Statistical significance of differences between controls and experimental samples was determined by two-sided, two-sample t-tests, for unequal standard deviations. In the figures, *** represents p≤0.001, ** represents p≤0.01, * is for p≤0.05, and ^###^=p≤0.001 represents decline from control value or from the ARSB silencing-induced value.

## 3.0 Results

### 3.1 Inhibition of tumor growth and improved survival following treatment with exogenous N-acetylgalactosamine-4-sulfatase (Arylsulfatase B; ARSB)

Cultured B16F10 cells demonstrated significant growth inhibition following treatment with rhARSB at doses ranging from 0.5-2.0 ng/ml for up to 24 hours (p<0.001, two-sided two sample t-test, unequal standard variation) (**Fig. 1A**). Following subcutaneous (SC) injection of 250,000 B16F10 cells (ATCC) in the right flank of 9-10-week-old female C57Bl/6J mice, control (n=12) and mice treated with rhARSB (n=28) were observed for up to 31 days. Doses of ARSB ranged from 0.1-0.4 mg/kg administered SC around the tumor. Treatment groups and the treatment schedules are presented (**Fig. 1B**), including one group with more frequent administration. The treatment groups are identified as groups 2, 3, 4, and 5, and the saline control is group 1. Treatment groups 3, 4 and 5 each had significant inhibition of tumor growth compared to saline control (p<0.0001, mixed effect model), with mean values presented (**Fig. 1C**). The lowest dose group (Group 2; 0.1 mg/kg) showed no significant inhibitory effect on tumor growth (p=0.069). No significant differences in inhibition effect occurred between the treatment groups 3, 4, and 5 (**Fig. 1C**). The p-values for the pairwise comparisons of tumor volume over the followup time between each pair of groups are presented (**Fig. 1D**). Representative tumor size by group is presented for Day 16 following tumor inoculation (**Fig. 1E**). Mean tumor volume on Day 16 post-inoculation in Groups 2-5 was 0.23 ± 0.15 cm^3^ (n=28), compared to 1.05 ± 0.7 cm^3^ in the controls (n=9) (p=0.006). Tumor weight of representative excised lesions at the time of spontaneous death or euthanasia shows significantly lower tumor weight in the treated mice in groups 2-5 (1.78 ± 0.65 g; n=18) compared to controls (3.19 ± 0.48 g; n=6; p=0.0001) (**Fig. 1F**).

The probability of survival up to day 21 was significantly increased in the treated mice, combining all treatment groups compared to the untreated control group (p=0.0391, log rank test) (**Fig. 1G**). Representation as the negative log of survival demonstrates improved survival in the treated mice at day 21, compared to untreated control (**Fig. 1H**). Six control mice died by day 18 post-tumor injection, in contrast to no deaths in treated mice by day 18. Three mice treated with ARSB had no palpable tumor when euthanized. No metastases were visible at the time of spontaneous death or euthanasia in any of the mice.

### 3.2 Declines in total sulfated glycosaminoglycans, chondroitin sulfate, and chondroitin 4-sulfate

In the untreated mice with tumors, total sulfated glycosaminoglycan content of the tumors was 24.11 ± 1.59 µg/mg protein, compared to 18.04 ± 1.06 µg/ml protein in treated mice (**Fig. 2A**). Total chondroitin sulfate tumor content was 14.25 ± 0.91 µg/mg protein in the control mice, compared to 9.43 ± 0.64 µg/mg protein (p=0.03, n=6 per group) in the mice treated with exogenous ARSB (**Fig. 2A**). This decrease is largely attributable to the decline in chondroitin 4-sulfate (C4S) from 8.73 ± 0.06 µg/ml to 4.20 ± 0.35 µg/ml (p<0.0001, n=6 per group) by ARSB treatment (**Fig. 2B**). Total sulfotransferase activity was reduced in the treated tumor tissue (p<0.001, n=6 per group) (**Fig. 2C**).

**FIG. 2.**
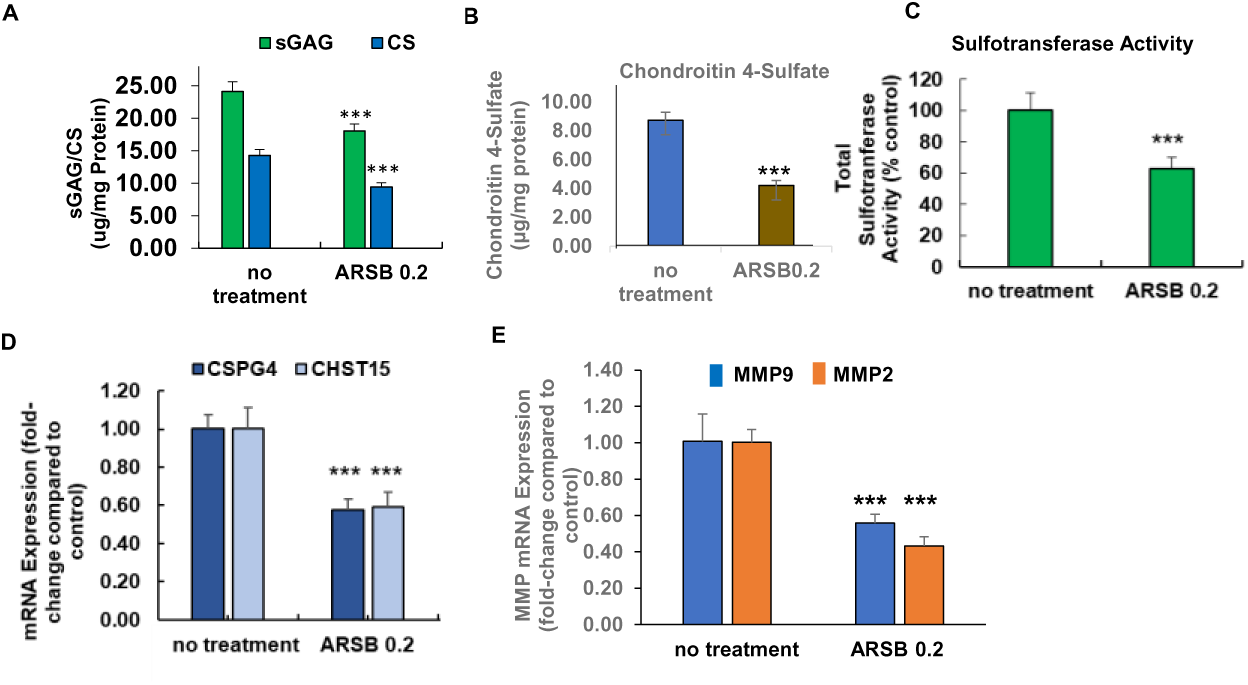
Effects of exogenous Arylsulfatase B on chondroitin 4-sulfate and associated parameters in B16F10 melanoma. **A.** Total chondroitin sulfate and total sulfated glycosaminoglycans declined significantly following treatment with exogenous ARSB (p<0.001, two sample t-test, two-tailed, unequal variance, n=6). **B.** Chondroitin 4-sulfate (C4S) was measured by the Blyscan assay following immunoprecipitation with C4S antibody and was markedly reduced in the tumor tissue from the treated mice (p<0.001, two sample t-test, two-tailed, unequal variance; n=6 for each group). **C.** Sulfotransferase activity was reduced in the tumor tissue from the treated mice compared to control (p<0.001, two sample t-test, two-tailed, unequal variance, n=6 for each group). **D.** mRNA expression of CSPG4 and CHST15 was reduced following exogenous ARSB (p<0.001, two sample t-test, two-tailed, unequal variance, n=6 for each group). **E.** mRNA expression of pro-MMP2 and MMP9 was reduced following ARSB treatment (p<0.001, two sample t-test, two-tailed, unequal variance, n=6 for each group). [ARSB=arylsulfatase B=N-acetylgalactosamine-4-sulfatase; con=control; consi=control siRNA; CHST15=carbohydrate sulfotransferase 15= chondroitin 4-sulfate 6-O-sulfotransferase; CS=chondroitin sulfate; CSPG4=chondroitin sulfate proteoglycan; gal3si=galectin-3 siRNA rh=recombinant human ARSB; sGAG=sulfated glycosaminoglycan; si-siRNA]

In the tumor tissues, mRNA expression of CSPG4 and CHST15 declined following treatment with exogenous ARSB (p<0.001, two-sample t-test, two-tailed, unequal variance, n=6) (**Fig. 2D**). mRNA expression of MMP9 and pro-MMP2 was also significantly reduced (p<0.001) (**Fig. 2E**).

### 3.3 Reduced binding of SHP2 to chondroitin 4-sulfate following exogenous ARSB leads to changes in activation of critical signaling molecules

Since previous cell-based experiments indicated that silencing of ARSB and the associated increase in chondroitin 4-sulfation enhanced the binding of SHP-2 (PTPN11) with C4S and reduced the SHP2 activity, the impact of exogenous ARSB on SHP2 in the tumor tissue was addressed. Treatment of the melanoma tumors with exogenous ARSB increased the active phospho-SHP2 (**Fig. 3A**), attributable to decline in SHP2 bound with C4S (**Fig.**). Consistent with the increase in phospho-SHP2, phospho-ERK1/2 (**Fig. 3C**) and phospho-p38-MAPK (**Fig. 3D**) declined in the tumor tissue. Pearson correlation r between phospho-ERK1/2 and phospho-p38 was 0.99 (**Fig. 3E**).

**FIG. 3.**
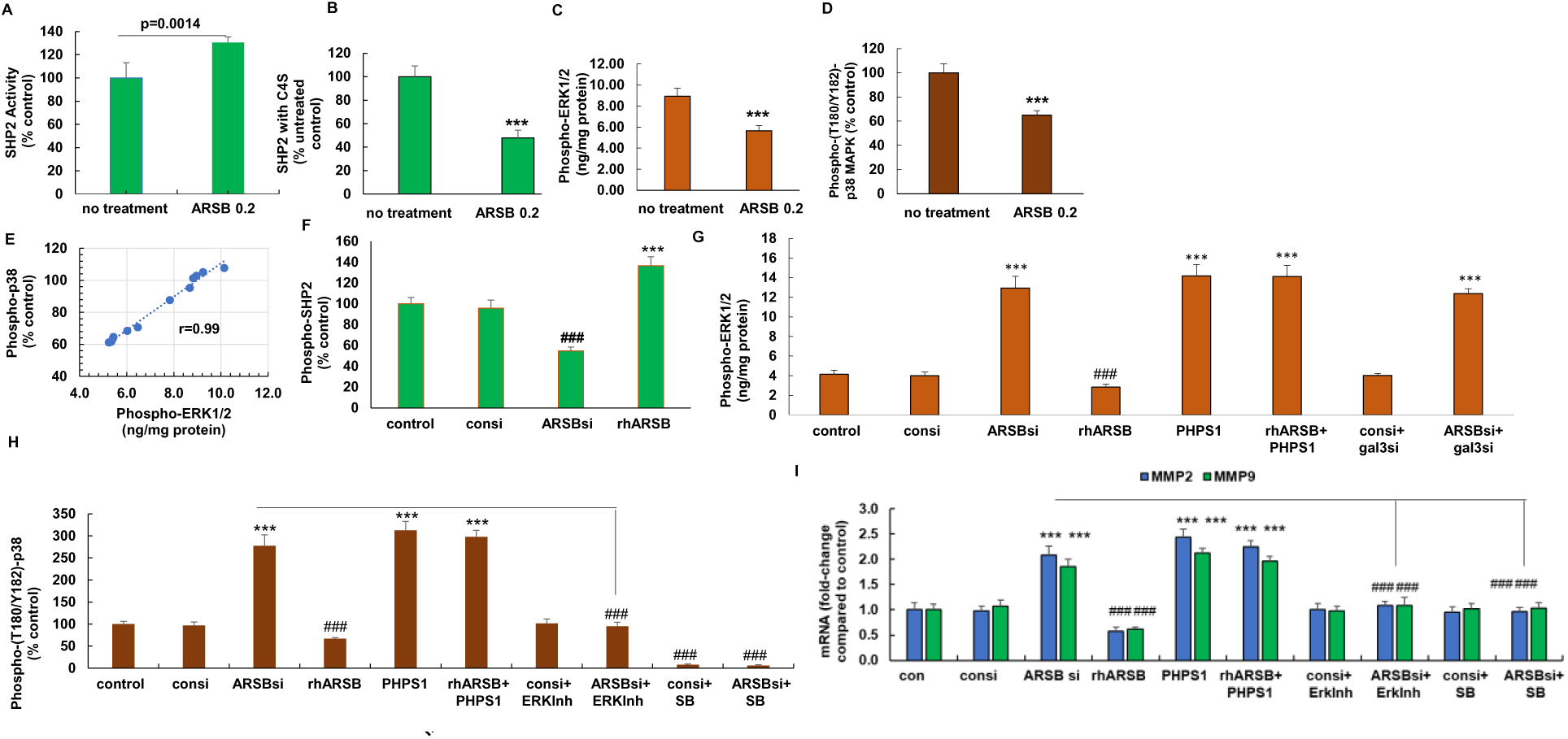
Signaling pathway leading to increased expression of MMPs following exogenous ARSB in B16F10 mouse melanoma and human A375 cells. **A.** SHP2 activity increased in the tumor tissue of the treated mice, compared to control (p=0.0014, two sample t-test, two-tailed, unequal variance; n=6 for each group). **B.** Binding of SHP2 with C4S declined in the treated tumor tissue, leading to the measured increase in SHP2 activity (p<0.001, two sample t-test, two-tailed, unequal variance; n=6 for each group). **C.** Following SHP2 activation, phospho-ERK1/2 was significantly reduced in the treated tumor tissue p<0.001, two sample t-test, two-tailed, unequal variance; n=6 for each group). **D.** Phospho-(Tyr180/Thr182)-p38MAPK was decreased following exposure to rhARSB in the treated tumor tissue, compared to untreated control (p<0.001, two sample t-test, two-tailed, unequal variance; n=6 for each group). **E.** Pearson correlation coefficient (r) between phospho-p38-MAPK and phospho-ERK1/2 in the tumor tissue was 0.99. **F.** In human A375 melanoma cells, SHP2 activity declined following ARSB silencing and increased following treatment by exogenous ARSB (p<0.001, two sample t-test, two-tailed, unequal variance, n=6 for each group). **G.** Consistent with the impact of SHP2 on phospho-ERK1/2, phospho-ERK1/2 declined following exogenous ARSB and increased when ARSB was silenced in the A375 cells. Galectin-3 silencing had no impact on the phospho-ERK1/2 (p<0.001, two sample t-test, two-tailed, unequal variance; n=6 for each group). **H.** Phospho-(T180/Y182)-p38 MAPK declined following exogenous ARSB and increased when ARSB was silenced (p<0.001, two sample t-test, two-tailed, unequal variance; n=6 for each group). Treatment with an inhibitor of ERK activation reversed the effect of ARSB silencing. SB203580 had the expected inhibitory effect. **I.** mRNA expression of pro-MMP2 and MMP9 in the A375 cells declined following exogenous ARSB (p<0.001, two sample t-test, two-tailed, unequal variance; n=6 for each group). Treatment with SB203580 or the ERK activation inhibitor blocked the ARSB siRNA-induced increases in MMP expression. [ARSBsi=ARSB siRNA; con=control; consi=control siRNA; ERKinh=ERK activation inhibitor; gal3si=galectin-3 siRNA; PHPS1=SHP2 inhibitor; rhARSB=recombinant human ARSB; SB=SB203580]

Experiments to further elucidate the underlying signaling pathways were performed in human A375 melanoma cells. Reciprocal effects of ARSB silencing and treatment with exogenous ARSB (1 ng/ml) were evident on phospho-SHP2 (**Fig. 3F**), phospho-ERK1/2 (**Fig. 3G**), phospho-p38 MAPK (**Fig. 3H**), and MMP9 and MMP2 expression (**Fig. 3I**). The signaling pathway, as elucidated by the effect of inhibitory molecules, indicates: 1) SHP2 inhibition by PHPS1 has similar effects as ARSB silencing and inhibits phospho-ERK1/2, phospho-p38 and expression of MMP-2 and MMP-9; 2) ERK activation inhibitor blocks phospho-p38 MAPK and reduces MMP2 and MMP9 expression; and 3) SB203580, the p38 MAPK inhibitor, blocks expression of MMP2 and MMP9.

### 3.4 Decline in free galectin-3 following exogenous ARSB leads to reduced expression of CSPG4 and CHST15

In the ARSB-treated tumor tissue, free galectin-3 and nuclear Sp1 declined, compared to the control (**Fig. 4A,**4B). In the cultured A375 human melanoma cells, silencing ARSB and exogenous ARSB had inverse effects. Following exogenous ARSB and the anticipated decline in chondroitin 4-sulfation, free galectin-3 and nuclear Sp1 were less (**Fig. 4C, 4D**). Galectin-3 silencing inhibited the ARSB siRNA-induced increase in nuclear Sp1 (**Fig. 4D**). Expression of CSPG4 declined following exogenous ARSB and increased following ARSB silencing (**Fig. 4E**). Galectin-3 silencing blocked the effect of ARSB silencing on CSPG4 expression, but PHPS1 and SB203580 had no inhibitory effect. Exposure to mithramycin, an inhibitor of Sp1 DNA binding, blocked the ARSB siRNA induced increase in CSPG4 expression.

**FIG. 4.**
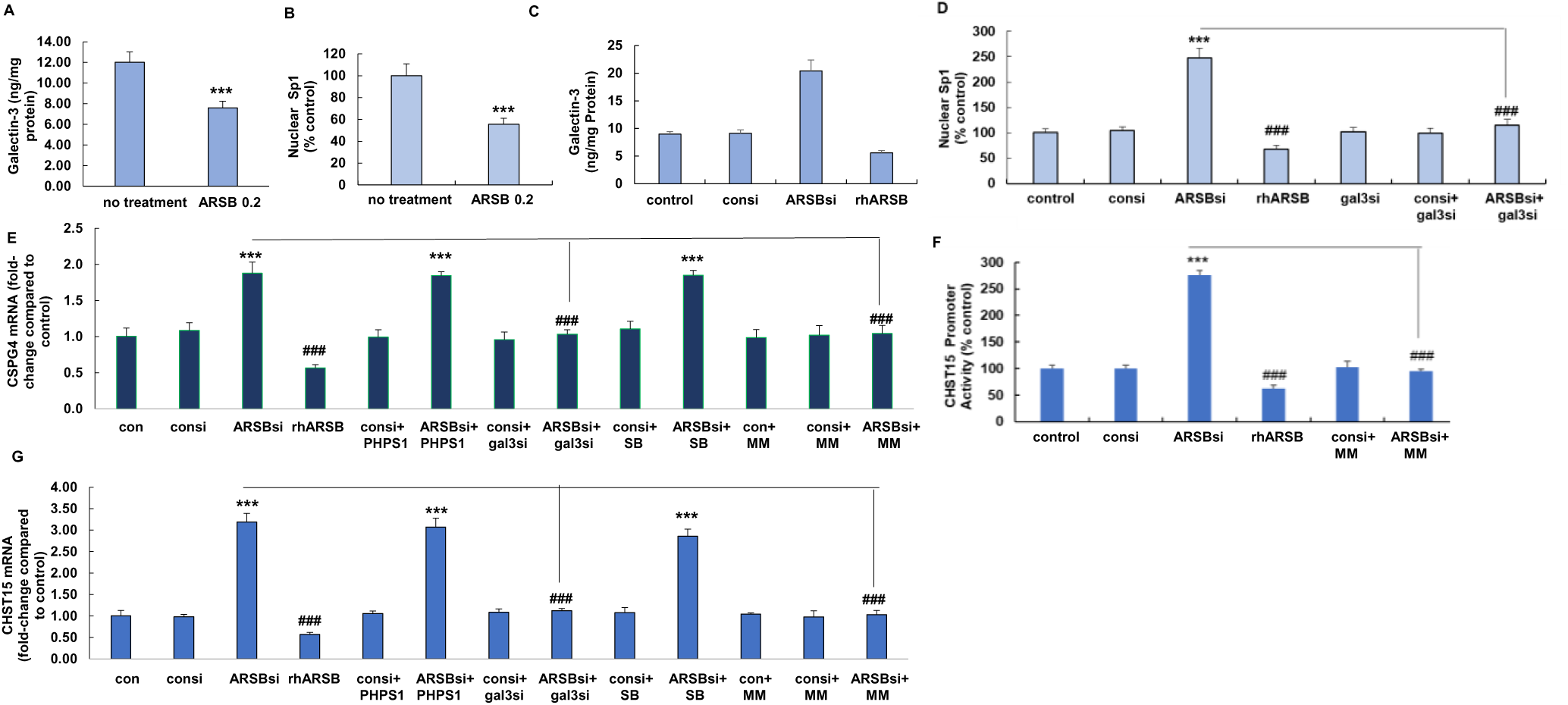
Galectin-3 and Sp1 mediate expression of CSPG4 and CHST15 initiated by modulation of ARSB in B16F10 mouse melanomas and human A375 cells. **A.** Free galectin-3 decreased following exogenous ARSB in the mouse melanomas (p<0.001, two sample t-test, two-tailed, unequal variance, n=6 in each group). **B.** Nuclear Sp1 in the melanoma tissue also declined following ARSB treatment (p<0.001, two sample t-test, two-tailed, unequal variance, n=6 in each group). **C.** In A375 cells treated with rhARSB, free galectin-3 declined. In contrast, ARSB silencing reduced the free galectin-3 (p<0.001, two sample t-test, two-tailed, unequal variance, n=6 in each group). **D.** Nuclear Sp1 declined when cells were treated with rhARSB. ARSB siRNA increased the nuclear Sp1, and this increase was inhibited by galectin-3 silencing (p<0.001, two sample t-test, two-tailed, unequal variance, n=6 in each group). **E.** mRNA expression of CSPG4 declined following treatment with rhARSB. In contrast, ARSB silencing increased expression, and the increase was inhibited by mithramycin and galectin-3 siRNA, but not by PHPS1 or by SB203580 (p<0.001, two sample t-test, two-tailed, unequal variance, n=6 in each group). **F.** CHST15 promoter activation increased following ARSB silencing and declined following treatment with exogenous ARSB (p<0.001, two sample t-test, two-tailed, unequal variance, n=6 in each group). The ARSB siRNA-mediated increase was inhibited by mithramycin. **G.** CHST15 expression increased when ARSB was silenced and declined following exogenous ARSB. The ARSB siRNA-induced increase was inhibited by galectin-3 siRNA and by mithramycin, but not by PHPS1 or SB203580 (p<0.001, two sample t-test, two-tailed, unequal variance, n=6 in each group). [ARSBsi=ARSB siRNA; CHST15 = N-acetylgalactosamine 4-sulfate 6-O-sulfotransferase; con=control; consi=control siRNA; CSPG4=chondroitin sulfate proteoglycan; ERKinh=ERK activation inhibitor-1; gal3si=galectin-3 siRNA; MM=mithramycin; rhARSB=recombinant human ARSB; PHPS1=SHP2 inhibitor; SB=SB203580]

Similar to the mechanism mediating CSPG4 expression, CHST15 promoter activation followed ARSB silencing and was reduced following exogenous ARSB (**Fig. 4F**). The ARSB siRNA-induced increase in CHST15 promoter activation was inhibited by treatment with mithramycin. CHST15 expression increased following ARSB silencing and was inhibited following exogenous ARSB (**Fig. 4G**). Treatment with PHPS1 or SB203580 had no effect on the ARSB siRNA-induced increase in CHST15, whereas galectin-3 siRNA and mithramycin inhibited the increase.

### 3.5 A schematic showing these signaling pathways is presented in **Fig. 5.** The findings in the mouse melanoma model are consistent with the pathways demonstrated in the A375 cells

**FIG. 5.**
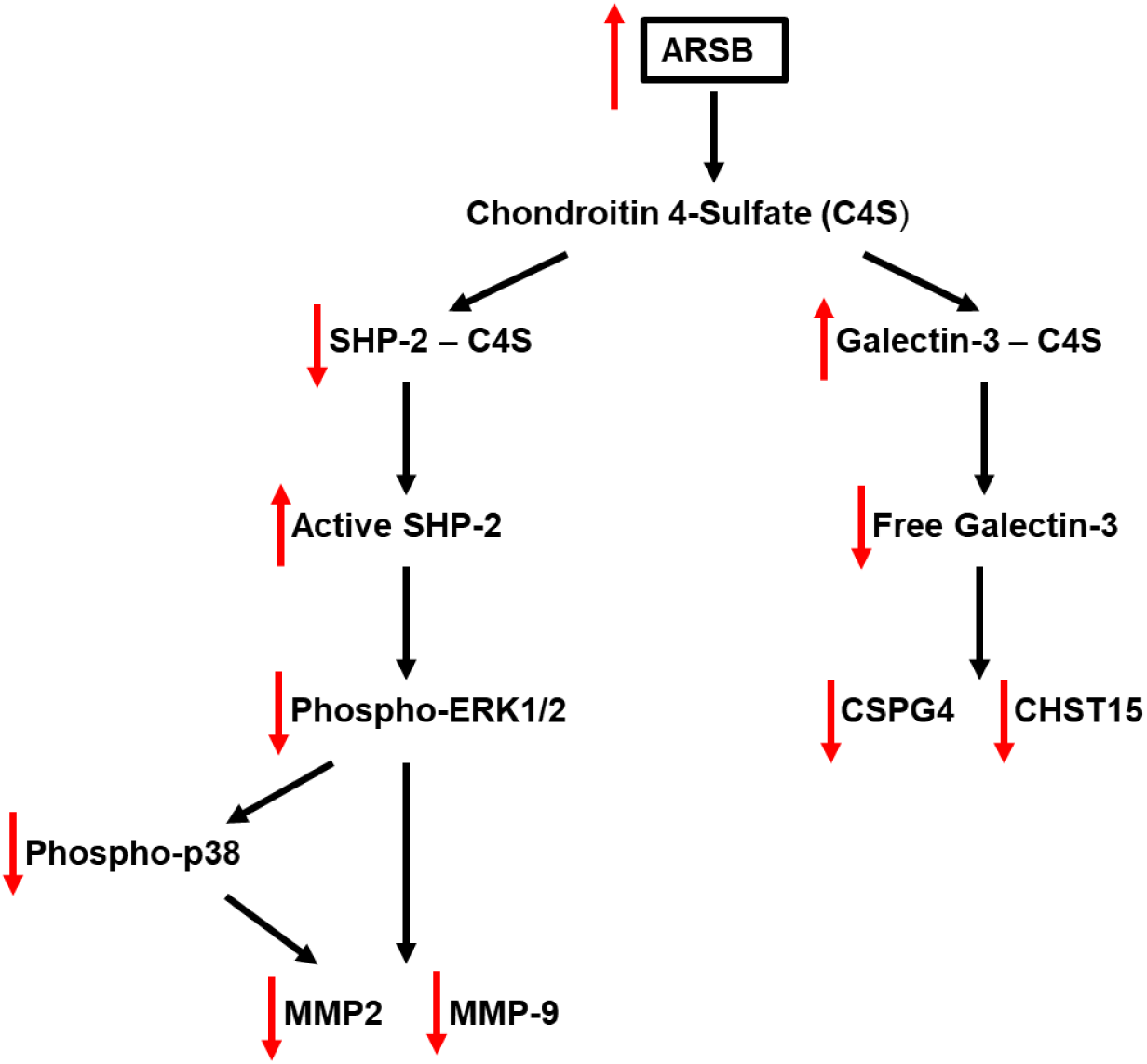
Schematic of signaling pathways in melanoma cells and tissue. This schematic displays the two pathways by which ARSB affects transcriptional events in the mouse melanoma tissue and the human A175 cell line. The distinct mechanistic pathways arise from differential binding with more or less highly sulfated C4S and lead to galectin-3/Sp1 mediated transcriptional effects and to effects on intracellular MAPK- phosphorylation signaling cascades requiring activation/inactivation of SHP2, ERK1/2, and p38 MAPK. [ARSB=Arylsulfatase B=N-acetylgalactosamine 4-sulfatase; C4S=chondroitin 4-sulfate; CHST15=N-acetylgalactosamine 4-sulfate 6-O-sulfotransferase; CSPG4=chondroitin sulfate proteoglycan; ERK=extracellular-regulated kinase; MMP=matrix metalloproteinase; SHP2=PTPN11=Tyrosine-protein phosphatase non-receptor type 11]

## 4.0 Discussion

Locally administered, exogenous ARSB markedly inhibited the growth of subcutaneous malignant melanomas in the B16F10 syngeneic mouse model in young, female C57Bl/6J mice. The recombinant ARSB, which is biologically active, reduced the content of C4S in the tumors. As demonstrated previously in cell-based experiments, the association of the critical molecules SHP2 and galectin-3 with C4S was modified by changes in ARSB [8,9]. Changes in availability of SHP2 and galectin-4 have crucial effects on cell signaling which impact on proliferation and survival.

Consistent with previous reports about the effect of decline in ARSB on SHP-2 and galectin-3, treatment with exogenous, bioactive ARSB increased the phospho-SHP2 activity and reduced phospho-ERK1/2 and phospho-p38 MAPK. Free galectin-3 was reduced following exogenous ARSB in the treated mice and A375 cells, attributable to enhanced binding with less sulfated C4S [9,34]. These changes affected transcriptional events, leading to declines in MMP2 and MMP9 and CSPG4 and CHST15.

Signaling pathways in the A375 cells show that the expression of the MMPs increased when ARSB was silenced and declined with exposure to exogenous ARSB. The increase in expression was blocked by the SHP2 inhibitor PHPS1, by the ERK activation inhibitor, and by SB203580, a specific inhibitor of p38-MAPK. In contrast to inhibition by PHPS1, galectin-3 silencing had no effect on ERK1/2 phosphorylation. PHPS1, ERK activation inhibitor peptide-1, and SB203580 blocked the increases in phospho-p38 and in MMP9 and pro-MMP2 expression which followed ARSB silencing.

In contrast to the SHP2-mediated transcriptional effects, neither PHPS1, nor SB203580, blocked the ARSB-silencing-induced increases in expression of CSPG4 or CHST15. Expression of CSPG4 and CHST15 was inhibited by galectin-3 silencing and by mithramycin, which blocks Sp1 DNA binding.

The cell-based results corroborate the measurements from the mouse melanomas and imply that the decline in tumor growth, and the resulting improvement in survival, are mediated by the impact of exogenous ARSB on chondroitin 4-sulfation. Other transcriptional mechanisms and metabolic changes in the tumor cells are likely also occurring and affecting the tumor growth. The specific effects of decline in CSPG4 and CHST15 on the interactions of tumor cells with neighboring cells, inflammatory cells, and the extracellular matrix require further evaluation. This is the first report of the potential association of increased CHST15 expression in melanoma, although the contribution of CHST15 in malignancy is increasingly recognized [35]. It is possible that a decline in abundance of CSPG4 and CHST15 by treatment with exogenous ARSB may improve recognition of the tumor cells by infiltrating immune cells or may enhance contact inhibition.

The mechanisms considered in this report control vital pathways, including phospho-ERK-phospho-38 MAPK activation and galectin-3-mediated transcriptional events. Other work has shown a vital role for galectin-3 in regulation of insulin receptor responsiveness [36,37], and suggests that treatment with ARSB and enhanced binding of galectin-3 with less sulfated C4S may lessen insulin resistance. Downstream signaling effects may thereby modulate phospho-AKT and other vital metabolic pathways.

The impact of sulfation on phosphorylation poses a new approach to address malignant transformation and new opportunities for intervention. For these experiments, bioactive, exogenous rhARSB was used and is anticipated to enter cells through the mannose-6 receptor, similar to the uptake pathway in treatment of MPS VI, and does not require post-translational modification by the formylglycine-generating enzyme [38–40]. Potential use of ARSB gene transduction in tumors poses questions about effectiveness of expression, post-translational modification, and intracellular transport and localization of ARSB which require further examination. The use of ARSB gene replacement in MPSVI provides insight into potential genetic approaches to treatment with ARSB which may be relevant to future initiatives with ARSB in malignancy.

### 5.0 Conclusion

The findings presented in this report demonstrate effectiveness of exogenous ARSB on inhibition of the progression of subcutaneous mouse melanomas and elucidate mechanisms which lead to declines in matrix metalloproteinases 2 and 9 and chondroitin sulfate proteoglycan 4, known biomarkers of melanoma progression. Attention to ARSB and chondroitin 4-sulfate-mediated regulation of transcriptional events provides an innovative approach to address the underlying molecular basis of melanoma progression.

## Acknowledgments

Research reported in this publication was supported in part by the University of Illinois Cancer Center Biostatistics Shared Resource (BSR) core (ZC and JT). The facilities of the VA and VA Merit Review grant to JKT supported the research. The content is solely the responsibility of the authors and does not necessarily represent the official views of the National Institutes of Health or the VA.

## Author Contributions

SB: Conceptualization, Data curation, Formal analysis; Investigation; Writing original draft

IO: Conceptualization, Investigation, Methodology

JT: Methodology, Software, Writing original draft

ZC: Funding acquisition, Methodology, Software, Supervision, Writing-original draft and review and editing

JKT: Conceptualization, Funding acquisition, Investigation, Methodology, Project Administration, Resources, Supervision, Validation, Writing-original draft and review and editing

## Data Availability

Data are available by communication with JKT.

## Notes

**Conflict of Interest:** The authors have no conflict of interest with this report.

### Competing Interest Statement

The authors have declared no competing interest.

